# Local read haplotagging enables accurate long-read small variant calling

**DOI:** 10.1101/2023.09.07.556731

**Authors:** Alexey Kolesnikov, Daniel Cook, Maria Nattestad, Brandy McNulty, John Gorzynski, Sneha Goenka, Euan A. Ashley, Miten Jain, Karen H. Miga, Benedict Paten, Pi-Chuan Chang, Andrew Carroll, Kishwar Shafin

## Abstract

Long-read sequencing technology has enabled variant detection in difficult-to-map regions of the genome and enabled rapid genetic diagnosis in clinical settings. Rapidly evolving third-generation sequencing platforms like Pacific Biosciences (PacBio) and Oxford nanopore technologies (ONT) are introducing newer platforms and data types. It has been demonstrated that variant calling methods based on deep neural networks can use local haplotyping information with long-reads to improve the genotyping accuracy. However, using local haplotype information creates an overhead as variant calling needs to be performed multiple times which ultimately makes it difficult to extend to new data types and platforms as they get introduced. In this work, we have developed a local haplotype approximate method that enables state-of-the-art variant calling performance with multiple sequencing platforms including PacBio Revio system, ONT R10.4 simplex and duplex data. This addition of local haplotype approximation makes DeepVariant a universal variant calling solution for long-read sequencing platforms.

## Introduction

Long-read sequencing technology can reach low-mappability regions where short-reads have difficulty in mapping correctly [1, 2, 3, 4, 5]. Long-read sequencing has helped to generate highly contiguous genome assemblies [6, 7, 8], extend variant benchmarking to complex regions [9, 10] and has improved the quality and completeness of reference genomes [11, 12, 13, 14].

Variant detection with long-reads takes advantage of the high mappability of long reads to reach difficult-to-map regions of the genome [1, 15, 16]. Long-read sequencing technology have lower base-level accuracy compared to short read technologies [1, 17, 18, 19, 20]. However, variant callers based on deep neural networks (DNN) can produce highly accurate variant calls with long-reads [16, 21, 22]. Recently, high-throughput long-read sequencing paired with accurate variant calling has demonstrated fastest genetic diagnosis in clinical setting [23, 24, 25]. Also accurate long-reads shows higher diagnosis rate among individuals who were previously undiagnosed [26, 27]. Long-read sequencing paired with methods based on machine learning for variant detection shows promise to have a far reaching impact in healthcare [28].

DNN-based variant callers not only take advantage of the mappability of the long reads, it also uses local haplotyping information to inform the neural network for accurate genotyping [16, 21]. Previously [29], we have shown that using local haplotype information to determine the genotype of a variant improves the accuracy of variant detection with PacBio long-reads. In this three-step approach, a first round of variant detection is performed using DeepVariant [30] without haplotag information in the reads. Then, WhatsHap [31] is used to haplotag the reads. Finally, we run DeepVariant with the haplotag information to produce higher quality variants. A similar approach was taken for PEPPER-Margin-DeepVariant [16] pipeline for nanopore long-reads where Margin is used to haplotag the reads.

Other Long-read germline variant callers sunch as Clair3 [21] and Medaka [32] use local haplotagging information from haplotagging methods like WhatsHap [31], Margin [33] or LongPhase [34]. Although the previously described three-step approach for variant detection provides accurate variants, it is difficult to tune three separate methods for each new datatype or platform introduced for long-reads. The long-read platforms like Pacific Biosciences (PacBio), and Oxford Nanopore Technologies (ONT) are rapidly evolving [1, 35, 36, 37] and a simpler and universal variant calling approach that can enable accurate variant calling across different platforms is desirable.

Pacific Biosciences (PacBio) is a single-molecule real-time (SMRT) sequencing platform that uses circular consensus sequencing [29]. Recently, PacBio introduced a high-throughput machine called Revio [38] that uses transformer-based consensus sequence correction method DeepConsensus [39] on the instrument to generate highly accurate (99.9%) reads with length between 15kb and 20kb. The PacBio Revio machine can generate 30x human genome with one SMRT cell compared to 8x per SMRT cell with the previous Sequel-II machine which lowers the overall cost and turnaround time [40, 41, 42].

Oxford Nanopore Technologies (ONT) introduced R10.4 chemistry with two read types simplex and duplex [43]. Compared to R9.4.1 chemistry, R10.4 provides better resolution for homopolymer detection [43, 44]. The simplex sequencing mode shows average read quality of 98% and duplex reads with 99% accuracy [44]. The nanopore long reads can be 100kb+ which makes it suitable for high-quality assemblies [7].

In this work, we introduce an approximate haplotagging method that can locally haplotag long reads without having to generate variant calls. Our approach uses local candidates to haplotag the reads and then the deep neural network model uses the haplotag approximation to generate high-quality variants. This approach eliminates the requirement for having the first two steps for haplotagging the reads and reduces the overhead for extending support to newer platforms and chemistries. We show that approximate haplotagging with candidate variants has comparable accuracy to haplotagging with WhatsHap. We report comparable or higher variant calling accuracy compared to DeepVariant-WhatsHap-DeepVariant approach with PacBio HiFi data. We demonstrate that extending support to the PacBio Revio machine can achieve high INDEL and SNP accuracy (SNP F1-score of 0.9993 and INDEL F1-score of 0.993). We also report support for ONT R10.4 simplex and duplex dataset with requiring PEPPER-Margin upstream of DeepVariant. We demonstrate INDEL F1-socre of 0.84 and SNP F1-score of 0.9976 for nanopore simplex data and INDEL F1-socre of 0.90 and SNP F1-score of 0.999 for duplex data which is to our knowledge the highest variant calling accuracy achieved with nanopore long-reads.

## Results

### Local haplotype approximation method for genotyping with DeepVariant

DeepVariant performs variant calling in three stages: `make_examples, call_variants` and `postprocess_variants`. Previously, we have described each stage in detail [30, 29, 16]. Here, we briefly summarize each stage:

- `make_examples:` This stage identifies candidate variants and creates a multichannel tensor representation (also known as an “example”) for each candidate based on the read pileup at the candidate position. Eachexample encodes features such as read bases, read quality, mapping quality, read support for candidate, and read strand.
- `call_variants:` Examples are then given to a Convolutional Neural Network (CNN) that determines the genotype likelihood for the candidate variant represented in each example.
- `postprocess_variants:` Finally, we take the likelihoods from the CNN and report variants with their assigned genotype.

Achieving accurate variant calling using long-read sequencing data requires calling variants twice often in a three-step process [16, 29] ((Figure 1a, DeepVariant-WhatsHap-DeepVariant):

**Figure 1.**
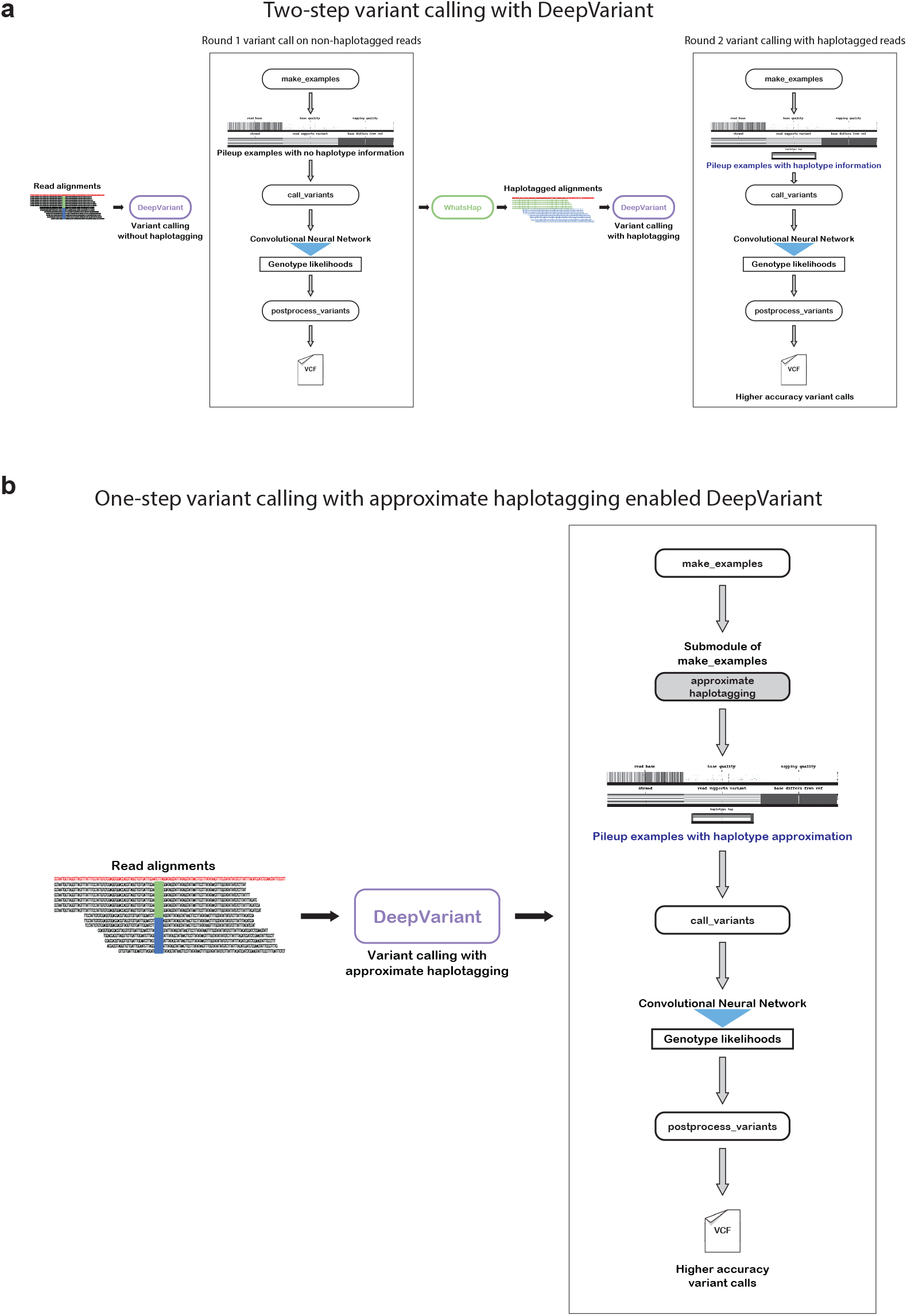
Illustration of different long-read variant calling schema. **a**. Two-step variant calling process that uses WhatsHap to haplotag the reads. **b**. Simplified variant calling process schema with approximate haplotagging method implemented within DeepVariant.

- **First round variant calling**: First, we run DeepVariant ont non-haplotagged alignment file to derive a set of variants.
- **Haplotagging**: Using the variants from the non-haplotagged alignment file a secondary method WhatsHap would be applied to haplotag the reads.
- **Second round variant calling**: Finally, the haplotagged read would be given to DeepVariant where DeepVariant would use extra features as haplotype channel to generate higher-quality variant calls.

The multi-step variant calling process of DeepVariant-WhatsHap-DeepVariant (Figure 1a) ntroduces overhead to extend support to newer long-read data types as multiple steps need to be optimized. In this work, we implement an approximate local haplotagging method within DeepVariant to locally haplotag the reads to avoid having to run an external haplotagging method or to have to reperform variant calling following initial haplotagging (Figure 1b). A description of the local haplotagging method is described here.

At each genomic position with a putative heterozygous variant, approximate haplotagging algorithm uses dynamic programming to calculate best scores for each possible phase assignment. Set of reads that overlap previous putative heterozygous variant location and target location are used to calculate the score. The best allele phasing can be calculated by backtracking from the best score for the last variant position. Once all alleles are assigned phases, read haplotags are assigned based on the majority of alleles that the read overlaps.

We use a graph to store read support information where each vertex represents an allele (alternate allele or ref). Edges are created to connect consecutive vertices if there is a read overlapping these alleles. The implementation of this method can be found in: https://github.com/google/deepvariant/blob/r1.5/deepvariant/direct_phasing.cc#L481C24-L481C24.

For each vertex, we calculate the score for phase assignment by adding the score from the previous vertices plus the number of reads that support that partition and overlap both the previous and current vertices. If a read starts after the previous position and overlaps the current vertices, it is added to the score. The implementation of this method can be found in: https://github.com/google/deepvariant/blob/r1.5/deepvariant/direct_phasing.cc#L282. During scoring, only alleles that have incoming edges are considered. When one of the vertices at a genomic position doesn’t have incoming edges the artificial edges are created fully connecting the vertex with all previous vertices. If there are no incoming edges for all vertices the phasing cannot be extended and we reinitialize the scores.

Here is a brief step-by-step description of the algorithm:

- Graph is built where each vertex is an allele (alt or ref). Edges are created to connect consecutive vertices if there is a read overlapping these alleles.
- Initial scores are calculated for all possible phasing for the first genomic position.
- Scores are calculated at each position using best scores calculated for previous vertices.
- The best score is backtracked from the last position. Each pair of alleles are assigned phases.
- Reads are assigned haplotags based on a set of alleles reads overlap.

The outcome of the local approximate haplotagging is a set of reads with haplotype association of haplotype-1, haplotype-2 or non-haplotagged reads for each 25kb chunk. Then for each candidate variant observed in the chunk, we generate examples through the make examples step of DeepVariant that creates a multi-channel representation of the pileup surrounding the candidate variant. The haplotype information is encoded in the haplotype channel and reads are also sorted based on the haplotype association of reads. The examples are provided to a Convolutional Neural Network (CNN) to provide genotype likelihood of homozygous to reference, heterozygous or homozygous alternate. Finally, in postprocessing step we take the output from the CNN and report the variant with a genotype and a likelihood of the genotype in Variant Call Format (VCF) file. A detailed description of the haplotagging algorithm with an example is provided in the online methods section.

### PacBio HiFi haplotype approximation and variant calling performance

DeepVariant haplotags long reads to improve genotyping accuracy during variant calling. We trained Deep-Variant models with no haplotagging information, haplotagging information from DeepVariant-WhatsHap bam, and the approximate haplotagging method. We trained the model on six Genome-In-A-Bottle (GIAB) samples (HG001, HG002, HG004, HG005, HG006, HG007) and kept HG003 as a hold out sample, we also holdout chromosme 20 as hold out. We then compared the variant calling performance on HG003 sample at different coverages.

In the variant calling performance analysis (Figure 2a and Supplementary table 1), we observe that DeepVariant model that uses no haplotagging information has lower variant calling performance at all coverages. The variant calling performance for SNPs with no haplotagging information can be considered comparable (F1-score at **15x**: 0.9960, **35x**: 0.9990) to the performance of variant calling with haplotagging information (F1-score at 15x: 0.996312, 35x: 0.999292). However, the INDEL performance of variant calling without haplotag information (F1-score at **15x**: 0.9529, **35x**: 0.9906) trails behind the performance of variant calling with haplotag information (F1-score at **15x**: 0.9701, **35x**: 0.9945) showing the importance of using haplotag information during variant calling with PacBio HiFi long reads.

**Figure 2.**
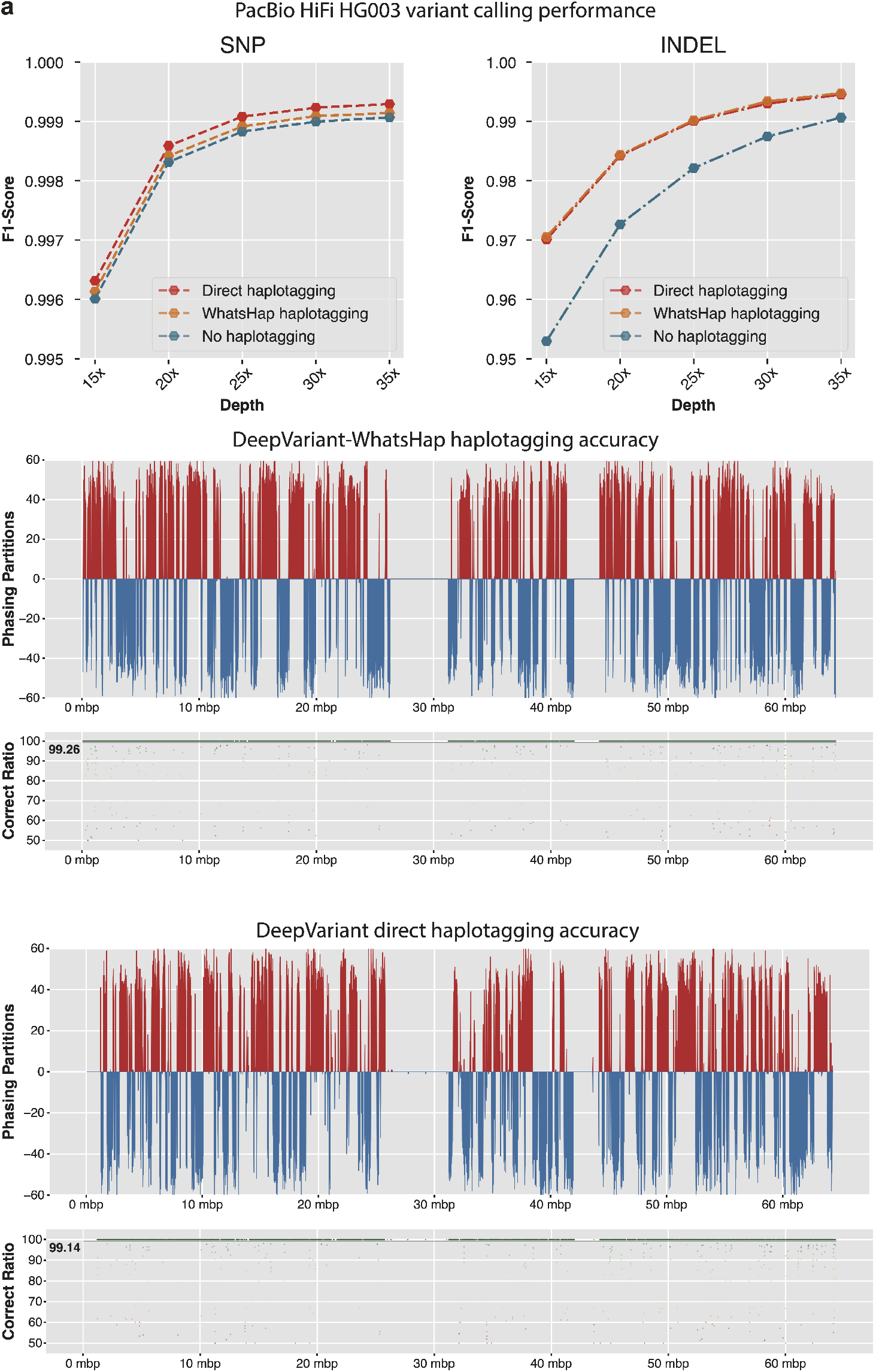
Effectiveness of local haplotagging approximation for variant calling. **a**. Variant calling accuracy of PacBio HiFi reads with no haplotagging, haplotagging with WhatsHap and approximate haplotagging. **b**. Haplotagging accuracy of WhatsHap and the local approximate haplotagging method.

We also observe that the variant calling performance of DeepVariant that uses approximate haplotagging built within the variant calling process is comparable to previously used WhatsHap-based pipeline. We see the SNP performance at each coverage, model trained with haplotag information from approximate haplotagging (F1-score **15x**:0.9963, **35x**: 0.9992) outperforms the WhatsHap-based model (F1-score **15x**: 0.9961, **35x**: 0.9947). For INDEL performance, we see that the performance of the model trained with aproximate haplotagging (F1-score **15x**: 0.9701, **35x**: 0.9945) is comparable to WhatsHap-based model (F1-score **15x**: 0.9705, **35x**: 0.9947). Overall, we see that the model that operates with haplotag information from approximate haplotagging has comparable or better variant calling performance than WhatsHap-based method that requires running the entire variant calling process multiple times.

We compared the approximate haplotagging performance of DeepVariant against WhatsHap-based haplotagging accuracy on HG002 chr20. We took GIAB trio phased variant calls and haplotagged the PacBio HiFi reads using whatshap and used that as the truth set for haplotype association for reads. For whatshap-based haplotagging we used DeepVariant to initially identify variants from unphased bam and then used whatshap haplotag to assign haplotags to reads. For approximate haplotagging with DeepVariant, we reported the read haplotype association of each chunk and merged them. Finally, we compared the read haplotagging accuracy against GIAB-based haplotagging.

Figure 2b shows DeepVariant approximate haplotagging accuracy of 99.14 is comparable to WhatsHap-based haplotagging accuracy of 99.26. Notably, DeepVariant-based haplotagging is performed on 25kbp chunks and merged at the end whereas WhatsHap operates on entire chromsome with high-quality variants from DeepVariant. We also observe the phasing partitions are comparably consistent between WhatsHap-based haplotagging and DeepVariant approximate haplotagging. Overall, the approximate haplotagging method improves variant calling accuracy (Figure 2a) of DeepVariant by providing accurate local haplotagging information (Figure 2b) for accurate genotyping.

### PacBio Revio vs Sequel-II variant calling performance

PacBio has announced a new sequencing instrument called Revio. Revio is high-throughput than previous Sequel-II. We analyzed the variant calling performance of DeepVariant with approximate haplotagging on PacBio Revio and Sequel-II instruments. The model is trained on GIAB samples from both sequencing platforms and HG003 is held out from the training. We analyzed the performance of DeepVariant at different coverages at various coverage between 5x to 30x coverage.

In figure 3a, we see the read length distribution of Revio is much wider compared to Sequel-II method as the library preparation for Sequel-II used a more precise gel-based method for read size selection. The empirical QV of both Revio and Sequel-II data are comparable.

**Figure 3.**
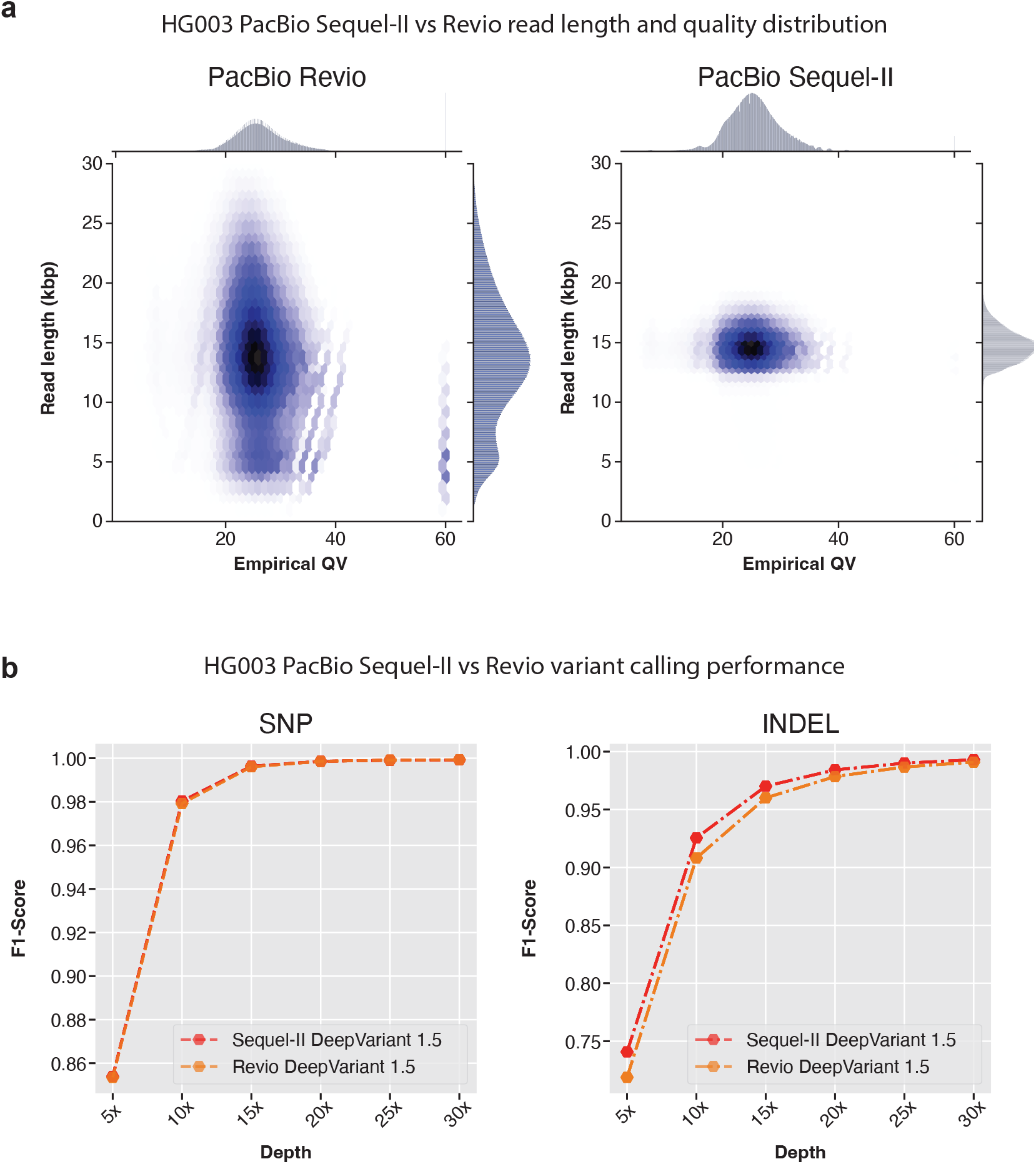
PacBio Revio and Sequl-II variant calling performance comparison. **a**. Read length distribution and empirical QV distribution of reads from Revio and Sequel-II. **b**. Variant calling performance of DeepVariant with Revio and Sequel-II data.

The variant calling performance of DeepVariant with Sequel-II and Revio are shown in figure 3b and supplementary table 2. Here we see the SNP calling performance of DeepVariant is comparable between Sequel-II and Revio platforms. The SNP variant calling performance (F1-scores of Sequel-II and Revio respectively, **5x**: 0.8539 vs 0.8535, **10x**: 0.9803 vs 0.9793, **15x**: 0.9963 vs 0.9960, **20x**: 0.9985 vs 0.9985, **25x**: 0.9990 vs 0.9991, **30x**: 0.9992 vs 0.9993) are very similar between two platforms at any given coverage. The INDEL performance of DeepVariant at different coverages show some difference (F1-scores of Sequel-II and Revio respectively, **5x**: 0.740798 vs 0.718782, **10x**: 0.92547 vs 0.908092, **15x**: 0.970101 vs 0.960039, **20x**: 0.984227 vs 0.978443, **25x**: 0.990044 vs 0.986717, **30x**: 0.993007 vs 0.990944). We observed the INDEL variant calling performance of DeepVariant with Sequel-II is higher compared to Revio at coverages between 5x to 25x. Whereas, at the suggested coverage of 30x, the variant calling performance with Revio is comparable to Sequel-II. The difference in the variant calling performance could be attributed to the read length distribution difference between the two datasets. However, Revio is high-throughput and the overall resources required to generate 30x data is much fewer compared to Sequel-II which makes Revio more practical for large-scale studies.

### Oxford nanopore simplex variant calling performance

Oxford Nanopore Technologies (ONT) introduced an updated and more accurate chemistry known as R10.4 which improves on homopolymer errors in nanopore reads. The error-rate of ONT R9 data requires preprocessing methods PEPPER-Margin to find candidates and haplotag the reads before variant calling with DeepVariant. However, with the improved sequencing quality from ONT R10.4 chemistry and approximate haplotagging method in DeepVariant, we are now able to train a model for ONT R10.4 that can call variants directly with DeepVariant without requiring complex preprocessing of PEPPER-Margin. Similar to previous training schemes, we held out HG003 from training the model.

In figure 4b and and supplementary table 3, 4, we compared the variant calling performance of ONT R10.4 data with three available variant callers, PEPPER, Clair3 and DeepVariant. Here, PEPPER referes to the PEPPER-Margin-DeepVariant pipeline and DeepVariant refers to DeepVariant with approximate haplotagging. In our comparison, we obseve that DeepVariant with haplotagging has comparable SNP variant calling performance at all coverages (SNP F1-score, **15x**: DeepVariant: 0.9903, PEPPER: 0.9875, Clair3: 0.9880, **30x**: DeepVariant: 0.9968, PEPPER: 0.9962, Clair3: 0.9957, **65x**: DeepVariant: 0.9976, PEPPER: 0.9977, Clair3: 0.9976). On the other hand, the INDEL variant calling performance of DeepVariant is higher compared to Clair3 and PEPPER at high and low coverages (INDEL F1-score, **15x**: DeepVariant: 0.7121, PEPPER: 0.7013, Clair3: 0.6902, **30x**: DeepVariant: 0.8412, PEPPER: 0.8345, Clair3: 0.8400, **65x**: DeepVariant: 0.8976, PEPPER: 0.8830, Clair3: 0.8800). Overall, the DeepVariant with approximate haplotagging shows it can derive high-quality variants from ONT R10.4 sequencing data.

**Figure 4.**
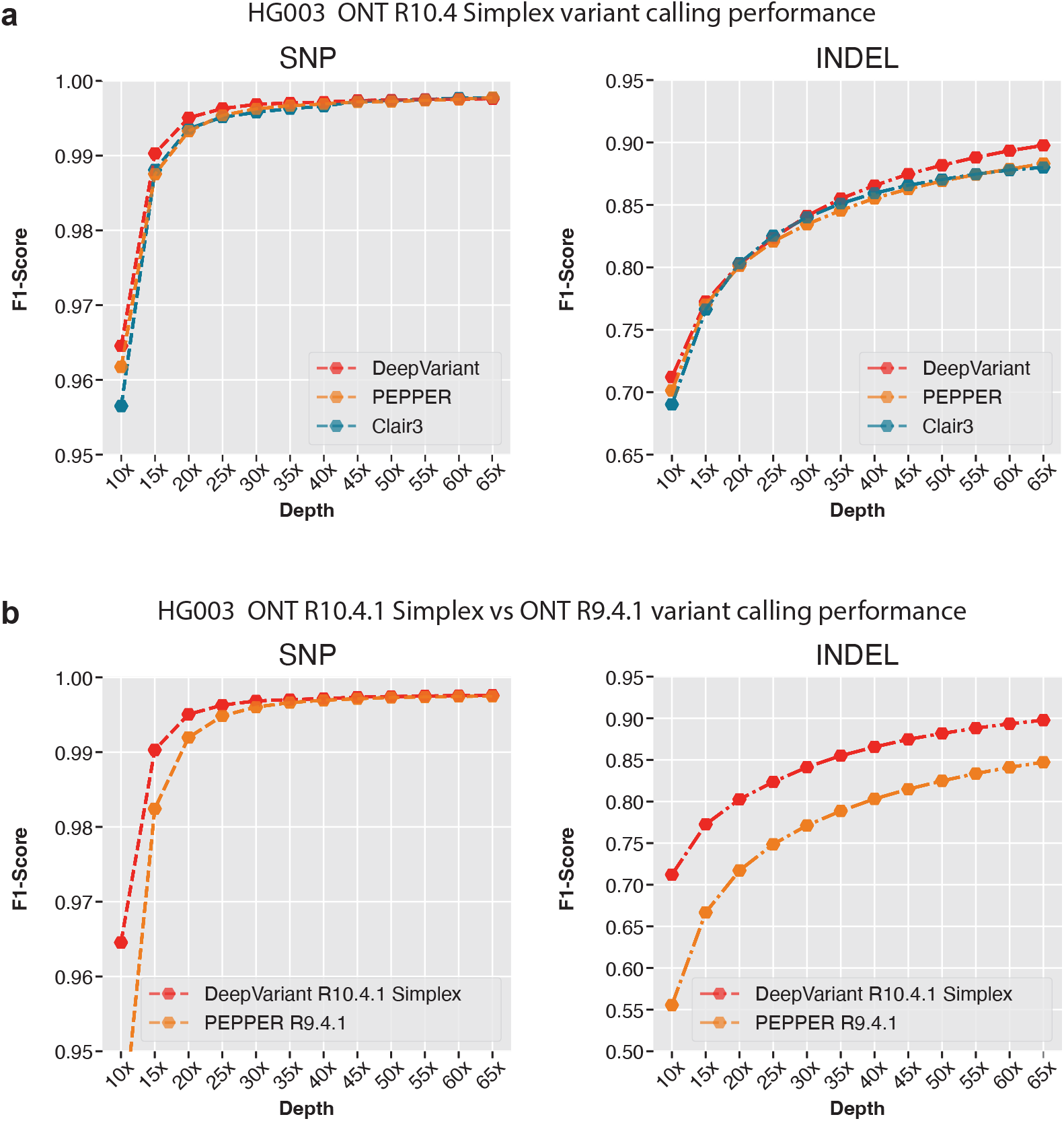
Variant calling performance comparison between Oxford nanopore R9.4.1 and R10.4 chemistry. **a**. Variant calling performance of R10.4 chemistry between DeepVariant, PEPPER and Clair3 variant callers. **b**. Variant calling performance of DeepVariant between R10.4 chemistry and R9.4.1 chemistry.

Figure 4b and supplementary table 5, 6 show the variant calling performance improvement between R9.4.1 and R10.4 chemistry. We observe that the SNP variant calling performance is comparable between R9 and R10 between 40x and 65x (SNP F1-score **40x**: R9: 0.9971, R10:0.9969,, 65x: R9: 0.9976, R10: 0.9974) coverage but R10 shows improvements between 10x and 35x coverage where the most improvement comes at lower coverage between 10x and 25x (SNP F1-score **10x**: R10: 0.9645, R9: 0.9371, **25x**: R10: 0.9962, R9: 0.9948) . The INDEL variant calling is highly improved with R10 chemistry at all coverages showing the improved data quality with the updated chemistry (INDEL F1-score **15x**: R10: 0.7725, R9: 0.6666, **30x**: R10: 0.8412, R9: 0.7712, **65x**: R9: 0.8472, R10: 0.8976).

### Improved variant calling with Oxford nanopore duplex sequencing

With the introduction of newer R10.4 chemistry, Oxford Nanopore also announced a new data type called duplex. We assessed the variant calling performance of DeepVariant on duplex and simplex data types of R10.4 chemistry. For this analysis we trained a model with R9.4 and R10.4 data where we had only HG002 available for duplex data. For each sample, we trained the model from chr1-chr19 and tuned on chr21-chr22 which left chr20 for evaluation. For evaluation, we compared the variant calling performance on chr20 of HG002 sample at different coverages.

Figure 5a shows the empirical QV vs read length distribution between simplex and duplex datasets. The simplex reads have a read length distribution that goes beyond 100kbp+ length reads. Although duplex read length is between 10kbp-50kbp, we observe major improvements in empirical QV with duplex data where the empirical QV reaches nearly Q30 whereas with simplex the empirical QV ranges between Q17 to Q20. With the improvements of empirical QV, duplex data is expected to deliver high-quality variant calls.

**Figure 5.**
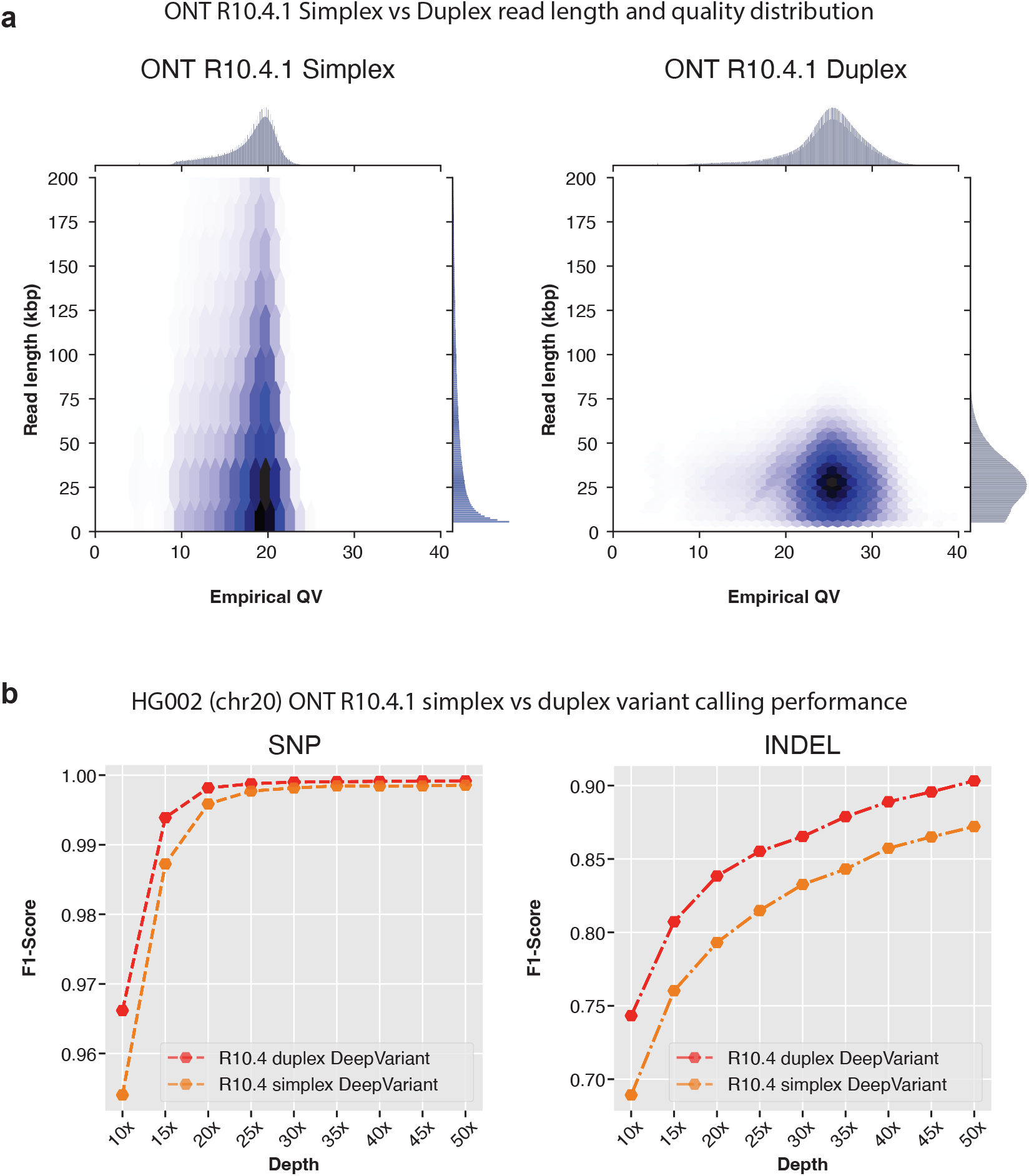
Comparison between Oxford Nanopore simplex and duplex datatypes with R10.4 chemistry. **a**. Read length distribution and empirical QV distribution of reads from simplex and duplex datatypes. **b**. Variant calling performance of DeepVariant with simplex and duplex datatypes.

The variant calling performance comparison between simplex and duplex datatype with DeepVariant shows duplex data improves variant calling performance for both SNP and INDEL (Figure 5b and supplementary table 7) at all coverages. The SNP variant calling performance at lower coverages with duplex data is improved compared to simplex data (SNP F1-score **10x**: Duplex: 0.9661, Simplex: 0.9540, **20x**: Duplex: 0.9981, Simplex: 0.9958, **30x**: Duplex: 0.9989, Simplex: 0.9981). At high-coverage between 40x-65x, simplex and duplex data shows comparable SNP variant calling performance (SNP F1-score **40x**: Duplex: 0.99911, Simplex: 0.9984, **50x**: Duplex: 0.9991, Simplex: 0.9985). For INDEL variant calling, duplex shows improvements at all coverages compared to simplex data (F1-score **10x**: Duplex: 0.7432, Simplex: 0.6892, **30x**: Duplex: 0.8652, Simplex: 0.8326, **50x**: Duplex: 0.9032, Simplex: 0.8720). Overall, the duplex datatype shows improvements in variant calling in all aspect making it an exciting new frontier for Nanopore sequencing.

## Discussion

The rapid improvement in long-read sequencing technology has shown it’s utility from high-quality genome assembly to rare disease diagnosis [45, 46, 11, 47]. Most variant calling methods that are tuned toward short-read sequencing technologies [48, 49] fail to derive and utilize the linkage information long-reads can provide during variant calling. Currently, most long-read variant callers require haplotagging to be done by an external method like WhatsHap [31] or Margin [16] which creates a complex multi-step variant calling process [29]. Besides inference, training and iteration on these models become difficult. In this work, we introduce an approximate local haplotagging method that can haplotag reads in 25kb chunks and improves the variant calling accuracy over non-haplotagged mode with PacBio HiFi data. We also extended the support to Oxford Nanopore Technologies R10.4 chemistry showing that it is possible to directly call variants from ONT data without multi-step PEPPER-Margin-DeepVariant setup.

The approximate haplotagging method we developed works efficiently on 25kb chunks of the genome and haplotags reads with comparable accuracy to WhatsHap. We show that the approximate haplotagging method achieves 99.14% correct haplotag ratio compared to 99.26% correct haplotag ratio of WhatsHap. We also show that models trained with haplotagging information from direct haplotagging method has comparable INDEL accuracy and higher SNP accuracy than models trained with haplotagging information with WhatsHap or with no-haplotagging information.

We show the variant calling performance on two PacBio platforms, Revio and Sequel-II. We show that the newer high-throuhput Revio’s variant calling performance with DeepVariant is comparable to Sequel-II platform at each coverage. At 30x coverage the performance of Sequel-II (F1-Score INDEL: 0.993, SNP: 0.9992) is comparable to Revio (F1-Score INDEL: 0.990, SNP: 0.9993). The performance difference at lower coverages between Revio and Sequel-II is suspected to be caused by the read length distribution difference between two datasets. We observed Sequel-II to have tighter band around 15kb read length whereas Revio had a wider read length distribution from 5kb to 25kb.

We extended DeepVariant to ONT R10.4 chemistry that has higher read-quality compared to R9.4.1 chemistry. We first show DeepVariant on R10.4 simplex data has the highest quality variant calls compared to PEPPER and Clair3. Then we show that DeepVariant with R10.4 is more accurate than PEPPER with R9.4 data. We show that with R10.4 chemistry it is possible to produce high-quality variants at lower coverages compared to R9 with demonstrable improvements in INDEL accuracy.

We also show that DeepVariant works seamlessly with R10.4 simplex and duplex data types. The duplex data has higher average read quality (average QV27) compared to simplex (average QV20) and produces even higher quality variant calls compared to simplex only data type. We show with R9.4.1 it is possible to achieve SNP F1-score of 0.999 at 30x Duplex coverage whereas 65x Simplex data achieves 0.997 SNP F1-score. For ONT INDEL performance, we see a noticeable improvement from R9.4.1 (INDEL F1-score 0.84), R10 simplex (INDEL F1-score 0.897) and R10 duplex (INDEL F1-Score 0.903). Although the INDEL improvements between platforms are on the positive side, we believe further iteration on the chemistry and data quality improvements would demonstrate further accuracy improvements in the future.

As we demonstrate a more generalized framework for long-read variant calling that uses haplotagging information, we believe future iterations of incorporating more long-read specific features would improve the variant calling accuracy further. For example, both PacBio and ONT can now produce methylation information [50, 51]. The newer methods can provide epigenetic profiles as well as canonical base calls [52, 53]. We believe incorporating methylation information in variant calling framework would further improve the accuracy.

## Acknowledgements

This study was supported by Google LLC (A.K., P.C., K.S., D.C., M.N., A.C.). B.P. was supported by the National Institutes of Health under award numbers R01HG010485, U24HG010262, U24HG011853, OT3HL142481, U01HG010961, and OT2OD033761. K.H.M. was supported by grants U01HG010971 and R01HG011274. We thank Trevor Pesout for the support of generating haplotag comparison plots. The content is solely the responsibility of the authors and does not necessarily represent the official views of Google LLC and the National Institutes of Health.

## Author Contributions

K.S., P.C., and A.C. conceived of the study and executed. A.K. refined and implemented approximate haplotagging algorithm. D.C. implemented haplotype channel, M.N. implemented alignment methods that improve long-read variant calling. K.S. performed the analysis and extended to ONT variant calling. A.C., P.C., A.K., D.C., M.N. and K.S. are core contributors and developers of DeepVariant. B.M., M.J. performed nanopore sequencing and initial analysis on the data quality. J.G. and S.G. provided analysis feedback. K.M., B.P. and E.A. provided guidance on the overall study and results. All authors approve of the final manuscript.

## Code Availability

Methods described here is publicly available through https://github.com/google/deepvariant where the haplotagging method can be found in https://github.com/google/deepvariant/blob/r1.5/deepvariant/direct_phasing.cc.

We have also made all input and output files publicly available. Please see the supplementary notes for access link.

## Online methods

### Approximate haplotagging

In this section, we describe the approximate haplotagging algorithm in details. We use graph data structure to perform haplotagging of reads. In a graph *G*, a set of vertices *V* represents alleles where a set of reads *R* overlaps these vertices. We need to assign haplotag of 1, 2 or 0 to all vertices so that this assignment has a maximum read support.

For a genomic position *n*, if we have two vertices *i* and *j* where *i* is assigned to phase-1 and *j* is assigned to phase-2, we define the best score as *S*(*V*_*n,i*_, *V*_*n,j*_). The set of supporting reads *R*(*V*_*n,i*_), *R*(*V*_*n,j*_) are stored for each possible phase assignment. Set of reads supporting the assignment *R*(*V*_*n,i*_) is calculated as a set of reads overlapping vertex *V*_*n,i*_ and preceding vertex *V*_*n*−1,*k*_ joined with a set of reads that overlap *V*_*n,i*_ and start after position *n* − 1.

At each step of the dynamic programming algorithm, we extend the best phase assignment calculated for the previous position. Final assignment is calculated by backtracking from the best score for the last genomic position. The two steps, initialization and recursion used are described below:

#### Initialization

The first genomic position of an interval (*n* = 1) and f or a ll p ossible p air of vertices *i* and *j*, we initialize the score as:

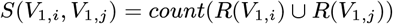

Initialization happens at the beginning of an interval or when the phasing cannot be extended from the previous position.

#### Recursion

During recursion we calculate scores at each position *n* for all vertex pairs (*V*_*n,i*_, *V*_*n,j*_) based on the previously observed vertex *V*_*n*−1,*k*_ and *V*_*n*−1,*l*_ where edges *E*(*V*_*n*−1,*k*_, *V*_*n,i*_) and *E*(*V*_*n*−1,*l*_, *V*_*n,j*_) exist in the graph connecting previous position *n* − 1 and current position *n*. The score is calculated as:

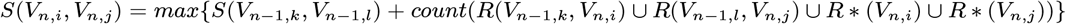

#### Backtracking and haplotag assignment

After scores are calculated for each vertex, the best scores are backtracked from the last position and each pair of alleles are assigned phases. Finally, reads are assigned haplotags based on a set of alleles reads overlap.

### Example of the approximate haplotagging algorithm

In figure 6, we illustrate a graph constructed for approximate haplotagging. In this graph each vertex represents an allele *A*_*n,m*_ where *n* is the genomic position and *m* is the allele. The alleles (*m*) are numbered for this demonstration. Set of reads overlapping each allele is shown next to it denoted as *read*1 to *read*11. The reads are placed as such it represents the alleles it supports in the graph.

**Figure 6.**
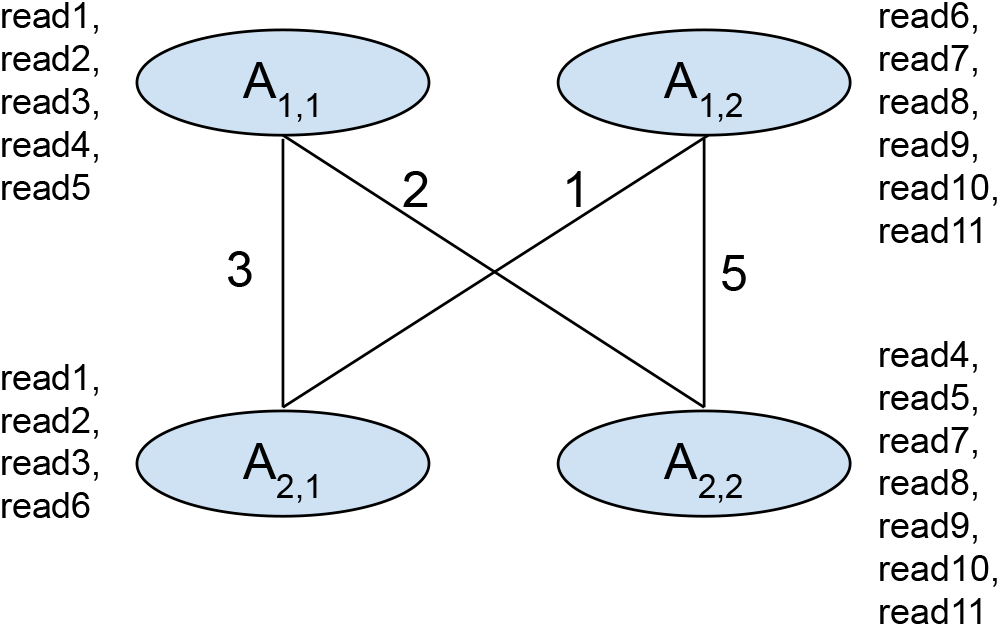
Illustration of a graph used to explain approximate haplotagging alogrithm.

In the first step, we initialize the scores for the genomic position *n* = 1:

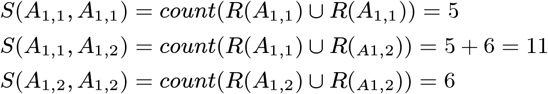

Then we calculate the scores of each pair of vertices based on the previously observed vertices:

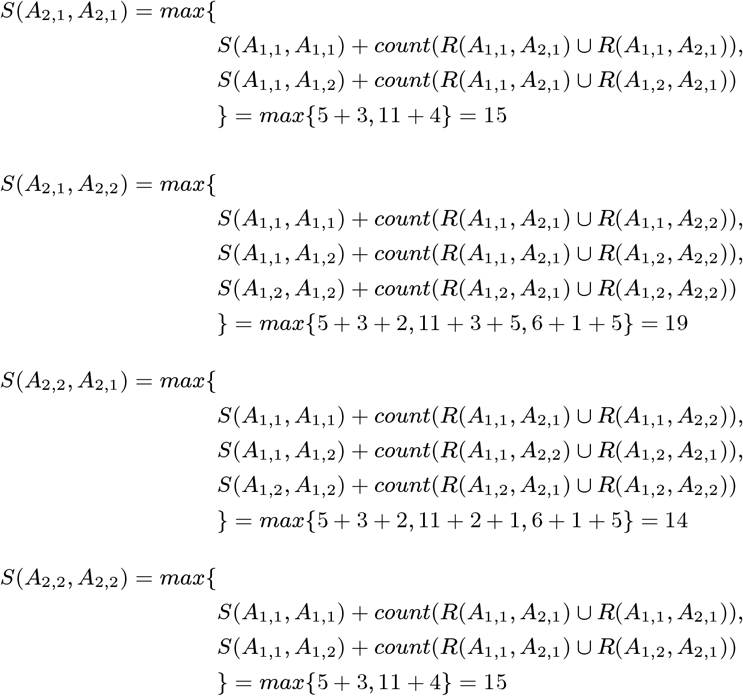

Then we calculate the best score at the last position. The best score for the last position is 19 where allele *A*_2,1_ is assigned to phase-1 and allele *A*_2,1_ is assigned to phase-2. Previous pair of alleles for this score is *A*_1,1_ with phase-1 and *A*_1,2_ with phase-2.

Using the best score, we assign phases to the vertices and haplotags to the reads. From the previous step we have alleles *A*_1,1_, *A*_2,1_ assigned to phase-1 and alleles *A*_2,1_, *A*_2,2_ assigned to phase-2. Reads: *read*1, *read*2, *read*3 overlap alleles *A*_1,1_, *A*_2,1_ both of which are phase-1. Therefore read1, read2, read3 are assigned haplotag-1.

Reads: *read*4, *read*5 overlap alleles *A*_1,1_, *A*_2,2_ are consecutively phase-1 and phase-2. In that case *read*4, and *read*5 cannot be phased. Following the same logic reads: *read*7, *read*8, *read*9, *read*10, *read*11 are assigned phase-2, and *read*6 cannot be phased.

Finally, these haplotag associations of the reads are used in the examples we generate for each candidate and the DNN model uses the information to generate accurate genotypes.

### Analysis methods

#### Read alignment and subsampling

We used `pbmm2` version 1.10 and `minimap2` [54] version `2.24-r1122` to align reads to the reference genome. We used `samtools` [55] version 1.15 for sampling alignment files at different coverages.

#### Variant calling and haplotagging

We used `PEPPER-Margin-DeepVariant` [16] version `r0.8, Clair3` [21] version `v1.0.0` for variant calling, for DeepVariant-WhatsHap-DeepVariant pipeline we used `v1.2.0` version of DeepVariant. For haplotagging with WhatsHap, we used WhatsHap version `v1.7`.

#### Benchmarking variant calls

For benchmarking variant calls, we used `hap.py` [56] version `v0.3.12`. The `hap.py` is available through `jmcdani20/hap.py:v0.3.12` in dockerhub. For we used GIAB v4.2.1 truth set [57] against `GRCh38` reference for all samples.

#### Haplotagging accuracy and natural switch determination

We used https://github.com/tpesout/genomics_scripts/haplotagging_stats.py to calculate the haplotagging accuracy [16].

#### Read accuracy estimation

We used `Best` [58] version `v0.1.0` for read accuracy analysis. For the analysis, we used GRCh38 as the reference to derive the empirical QV for each read.

