## Supplementary notes for "Local read haplotagging enables accurate long-read small variant calling"

#### Data availability

We have made all data including input BAMs, output VCF and analysis files publicly available: https://console.cloud.google.com/storage/browser/brain-genomics-public/publications/kolesnikov2023_dv_haplotagging/evaluation/

#### Commands used for analysis

##### pbmm2

We used `pbmm2` to align PacBio reads to the reference. Following is the command used:

```
docker run -it -v /data:/data \
quay.io/biocontainers/pbmm2:1.10.0--h9ee0642_0 \
pbmm2 align --preset HIFI --sort \
/data/Reference.fasta \
/data/inputs.fofn \
/data/output.pbmm2.bam
```

##### minimap2

We used `minimap2` to align ONT reads to the reference. Following is the command used:

```
minimap2 -k 17 -ax map-ont \
-t 95 REF.fasta INPUT.fastq.gz | samtools sort -@4 -m 4G > OUTPUT.bam
```

##### WhastHap

We used `WhastHap` to phase and haplotag PacBio and ONT data. Following is the command used:

```
whatshap phase --ignore-read-groups \
-o OUTPUT.phased.vcf -r REF.fasta \
INPUT.unphased.vcf INPUT.unhaplotagged.bam

whatshap haplotag \
--output OUTPUT.haplotagged.bam \
--reference /data/REF.fasta --ignore-read-groups \
OUTPUT.phased.vcf INPUT.unhaplotagged.bam
```

##### DeepVariant

We used `DeepVariant` variant caller to generate variant calls for PacBio and ONT data. Following is the command used:

```
docker run -v /data:/data \
google/deepvariant:1.5.0 \
/opt/deepvariant/bin/run_deepvariant \
--model_type=PACBIO \
--ref=/data/REF.fasta \
```

```
--reads=/data/INPUT.bam \
--output_vcf=/data/OUTPUT.vcf \
--output_gvcf=/data/OUTPUT.gvcf \
--num_shards=95 \
--logging_dir=/data/log_dir
```

We used `google/deepvariant:1.5.0` for analysis that includes approximate haplotagging and `google/deepvariant:1.2.0` for baseline comparison with no haplotagging and haplotagging with WhatsHap analysis. We used `-model_type=PACBIO` for variant calling on PacBio data and `-model_type=ONT_R104` for variant calling with ONT data.

#### Clair3

We used `Clair3` variant caller to generate variant calls for ONT data. Following is the command used:

```
sudo docker run -it -v /data:/data \
hkubal/clair3:latest \
/opt/bin/run_clair3.sh \
--bam_fn=/data/INPUT.bam \
--ref_fn=/data/REF.fasta \
--threads=95 \
--platform="ont" \
--model_path=/data/model_path/ \
--output=/data/output_dir/
```

We used `r1041_e82_400bps_sup_g615` model for variant calling R10.4 chemistry data with simplex and duplex types.

#### PEPPER

We used `PEPPER` variant caller to generate variant calls for ONT data. Following is the command used:

```
time docker run -it -v /data:/data \
kishwars/pepper_deepvariant:r0.8 \
run_pepper_margin_deepvariant call_variant \
-b /data/INPUT.bam \
-f /data/REF.fasta \
-o /data/output_dir/ \
-p output_prefix \
-t 95 \
--ont_r9_guppy5_sup
```

#### Hap.py

We used `hap.py` version `v0.3.12` to compare variant calls against GIAB truth set. Following is the command used:

```
docker run -it -v /data/:/data/ \
jmcdani20/hap.py:v0.3.12 /opt/hap.py/bin/hap.py \
/data/Benchmark.vcf.gz \
/data/INPUT.vcf.gz \
-f /data/Benchmark.bed \
-r /data/reference.fna \
-o /data/output/prefix \
--pass-only \
--engine=vcfeval \
--threads=95
```

#### BEST

We used best software `https://github.com/google/best` to assess the quality of the reads against reference genome.

```
./best \
INPUT.bam REF.fasta \
OUTPUT_DIR/OUTPUT_PREFIX \
--intervals-bed INTERVAL.bed -t 96
```

### Supplementary Tables

| Coverage | Mode | Type | Total | TP | FN | FP | Recall | Precision | F1-Score |
|---|---|---|---|---|---|---|---|---|---|
| 15x | No haplotag information | INDEL | 504501 | 475924 | 28577 | 19110 | 0.943356 | 0.962773 | 0.952966 |
| | | SNP | 3327495 | 3306618 | 20877 | 5629 | 0.993726 | 0.998302 | 0.996009 |
| | WhatsHap haplotagging | INDEL | 504501 | 487406 | 17095 | 12968 | 0.966115 | 0.975029 | 0.970551 |
| | | SNP | 3327495 | 3306574 | 20921 | 4750 | 0.993713 | 0.998567 | 0.996134 |
| | Approximate haplotagging | INDEL | 504501 | 486509 | 17992 | 12463 | 0.964337 | 0.975933 | 0.970101 |
| | | SNP | 3327495 | 3307835 | 19660 | 4831 | 0.994092 | 0.998543 | 0.996312 |
| 20x | No haplotag information | INDEL | 504501 | 488680 | 15821 | 12121 | 0.96864 | 0.976708 | 0.972658 |
| | | SNP | 3327495 | 3319839 | 7656 | 3549 | 0.997699 | 0.998933 | 0.998316 |
| | WhatsHap haplotagging | INDEL | 504501 | 496068 | 8433 | 7620 | 0.983284 | 0.985451 | 0.984366 |
| | | SNP | 3327495 | 3319757 | 7738 | 2778 | 0.997675 | 0.999165 | 0.998419 |
| | Approximate haplotagging | INDEL | 504501 | 495573 | 8928 | 7237 | 0.982303 | 0.986157 | 0.984227 |
| | | SNP | 3327495 | 3320920 | 6575 | 2814 | 0.998024 | 0.999154 | 0.998589 |
| 25x | No haplotag information | INDEL | 504501 | 494504 | 9997 | 8298 | 0.980184 | 0.984133 | 0.982155 |
| | | SNP | 3327495 | 3322610 | 4885 | 2911 | 0.998532 | 0.999125 | 0.998829 |
| | WhatsHap haplotagging | INDEL | 504501 | 499415 | 5086 | 4981 | 0.989919 | 0.99051 | 0.990214 |
| | | SNP | 3327495 | 3322526 | 4969 | 2245 | 0.998507 | 0.999325 | 0.998916 |
| | Approximate haplotagging | INDEL | 504501 | 499064 | 5437 | 4789 | 0.989223 | 0.990867 | 0.990044 |
| | | SNP | 3327495 | 3323614 | 3881 | 2231 | 0.998834 | 0.99933 | 0.999082 |
| 30x | No haplotag information | INDEL | 504501 | 497618 | 6883 | 5985 | 0.986357 | 0.988581 | 0.987468 |
| | | SNP | 3327495 | 3323336 | 4159 | 2521 | 0.99875 | 0.999243 | 0.998996 |
| | WhatsHap haplotagging | INDEL | 504501 | 501098 | 3403 | 3408 | 0.993255 | 0.993513 | 0.993384 |
| | | SNP | 3327495 | 3323326 | 4169 | 1889 | 0.998747 | 0.999432 | 0.99909 |
| | Approximate haplotagging | INDEL | 504501 | 500790 | 3711 | 3481 | 0.992644 | 0.99337 | 0.993007 |
| | | SNP | 3327495 | 3324370 | 3125 | 1989 | 0.999061 | 0.999403 | 0.999232 |
| 35x | No haplotag information | INDEL | 504501 | 499364 | 5137 | 4469 | 0.989818 | 0.991481 | 0.990649 |
| | | SNP | 3327495 | 3323559 | 3936 | 2257 | 0.998817 | 0.999322 | 0.999069 |
| | WhatsHap haplotagging | INDEL | 504501 | 501845 | 2656 | 2735 | 0.994735 | 0.994795 | 0.994765 |
| | | SNP | 3327495 | 3323619 | 3876 | 1830 | 0.998835 | 0.99945 | 0.999143 |
| | Approximate haplotagging | INDEL | 504501 | 501629 | 2872 | 2771 | 0.994307 | 0.994725 | 0.994516 |
| | | SNP | 3327495 | 3324633 | 2862 | 1852 | 0.99914 | 0.999444 | 0.999292 |

Supplementary Table 1: PacBio-HiFi variant calling performance of DeepVariant with no haplotagging, whatshap haplotagging and approximate haplotagging.

| Coverage | Platform | Type | Total | TP | FN | FP | Recall | Precision | F1_Score |
|---|---|---|---|---|---|---|---|---|---|
| 5x | Sequel II | INDEL | 504501 | 325065 | 179436 | 48949 | 0.64433 | 0.87124 | 0.740798 |
| | | SNP | 3327495 | 2568343 | 759152 | 119685 | 0.771855 | 0.955498 | 0.853914 |
| | Revio | INDEL | 504501 | 314909 | 189592 | 57900 | 0.624199 | 0.847147 | 0.718782 |
| | | SNP | 3327495 | 2565892 | 761603 | 119109 | 0.771118 | 0.955662 | 0.853529 |
| 10x | Sequel II | INDEL | 504501 | 454954 | 49547 | 24525 | 0.90179 | 0.950427 | 0.92547 |
| | | SNP | 3327495 | 3216400 | 111095 | 18094 | 0.966613 | 0.99441 | 0.980315 |
| | Revio | INDEL | 504501 | 445946 | 58555 | 32760 | 0.883935 | 0.933607 | 0.908092 |
| | | SNP | 3327495 | 3211531 | 115964 | 19959 | 0.96515 | 0.993828 | 0.979279 |
| 15x | Sequel II | INDEL | 504501 | 486509 | 17992 | 12463 | 0.964337 | 0.975933 | 0.970101 |
| | | SNP | 3327495 | 3307835 | 19660 | 4831 | 0.994092 | 0.998543 | 0.996312 |
| | Revio | INDEL | 504501 | 480852 | 23649 | 17008 | 0.953124 | 0.967055 | 0.960039 |
| | | SNP | 3327495 | 3306551 | 20944 | 5121 | 0.993706 | 0.998455 | 0.996075 |
| 20x | Sequel II | INDEL | 504501 | 495573 | 8928 | 7237 | 0.982303 | 0.986157 | 0.984227 |
| | | SNP | 3327495 | 3320920 | 6575 | 2814 | 0.998024 | 0.999154 | 0.998589 |
| | Revio | INDEL | 504501 | 492207 | 12294 | 9771 | 0.975631 | 0.981271 | 0.978443 |
| | | SNP | 3327495 | 3320738 | 6757 | 2801 | 0.997969 | 0.999158 | 0.998563 |
| 25x | Sequel II | INDEL | 504501 | 499064 | 5437 | 4789 | 0.989223 | 0.990867 | 0.990044 |
| | | SNP | 3327495 | 3323614 | 3881 | 2231 | 0.998834 | 0.99933 | 0.999082 |
| | Revio | INDEL | 504501 | 497096 | 7405 | 6223 | 0.985322 | 0.988116 | 0.986717 |
| | | SNP | 3327495 | 3323686 | 3809 | 2059 | 0.998855 | 0.999381 | 0.999118 |
| 30x | Sequel II | INDEL | 504501 | 500790 | 3711 | 3481 | 0.992644 | 0.99337 | 0.993007 |
| | | SNP | 3327495 | 3324370 | 3125 | 1989 | 0.999061 | 0.999403 | 0.999232 |
| | Revio | INDEL | 504501 | 499552 | 4949 | 4354 | 0.99019 | 0.991699 | 0.990944 |
| | | SNP | 3327495 | 3324580 | 2915 | 1727 | 0.999124 | 0.999481 | 0.999303 |

Supplementary Table 2: PacBio-HiFi variant calling performance comparison of DeepVariant on Sequel-II and Revio platforms at different coverages.

| Cov. | Caller | Type | Total | TP | FN | FP | Recall | Precision | F1 |
|---|---|---|---|---|---|---|---|---|---|
| 10x | DeepVariant | INDEL | 504501 | 316320 | 188181 | 69165 | 0.626996 | 0.824024 | 0.712133 |
| | | SNP | 3327495 | 3152000 | 175495 | 56238 | 0.947259 | 0.982476 | 0.964546 |
| | PEPPER | INDEL | 504501 | 298224 | 206277 | 48609 | 0.591127 | 0.862071 | 0.70134 |
| | | SNP | 3327495 | 3140268 | 187227 | 62700 | 0.943733 | 0.980428 | 0.961731 |
| | Clair3 | INDEL | 504501 | 334539 | 169962 | 132794 | 0.663109 | 0.719643 | 0.69022 |
| | | SNP | 3327495 | 3256496 | 70999 | 225280 | 0.978663 | 0.935315 | 0.956498 |
| 15x | DeepVariant | INDEL | 504501 | 359668 | 144833 | 69022 | 0.712918 | 0.843013 | 0.772527 |
| | | SNP | 3327495 | 3288775 | 38720 | 25744 | 0.988364 | 0.992235 | 0.990296 |
| | PEPPER | INDEL | 504501 | 350018 | 154483 | 55450 | 0.69379 | 0.865762 | 0.770294 |
| | | SNP | 3327495 | 3283461 | 44034 | 38814 | 0.986767 | 0.98832 | 0.987543 |
| | Clair3 | INDEL | 504501 | 365993 | 138508 | 86500 | 0.725455 | 0.812119 | 0.766345 |
| | | SNP | 3327495 | 3305838 | 21657 | 58142 | 0.993492 | 0.982723 | 0.988078 |
| 20x | DeepVariant | INDEL | 504501 | 380627 | 123874 | 65512 | 0.754462 | 0.857262 | 0.802584 |
| | | SNP | 3327495 | 3312923 | 14572 | 18308 | 0.995621 | 0.994506 | 0.995063 |
| | PEPPER | INDEL | 504501 | 374497 | 130004 | 57136 | 0.742312 | 0.870199 | 0.801184 |
| | | SNP | 3327495 | 3311580 | 15915 | 28778 | 0.995217 | 0.991387 | 0.993298 |
| | Clair3 | INDEL | 504501 | 382081 | 122420 | 66293 | 0.757344 | 0.854966 | 0.8032 |
| | | SNP | 3327495 | 3316247 | 11248 | 31521 | 0.99662 | 0.990588 | 0.993595 |
| 25x | DeepVariant | INDEL | 504501 | 395117 | 109384 | 62184 | 0.783184 | 0.868111 | 0.823463 |
| | | SNP | 3327495 | 3318240 | 9255 | 15473 | 0.997219 | 0.99536 | 0.996289 |
| | PEPPER | INDEL | 504501 | 389165 | 115336 | 55945 | 0.771386 | 0.876844 | 0.820741 |
| | | SNP | 3327495 | 3318979 | 8516 | 22297 | 0.997441 | 0.993329 | 0.99538 |
| | Clair3 | INDEL | 504501 | 393418 | 111083 | 56896 | 0.779816 | 0.876246 | 0.825223 |
| | | SNP | 3327495 | 3319621 | 7874 | 24503 | 0.997634 | 0.992676 | 0.995149 |
| 30x | DeepVariant | INDEL | 504501 | 407454 | 97047 | 58882 | 0.807638 | 0.877687 | 0.841207 |
| | | SNP | 3327495 | 3320004 | 7491 | 13466 | 0.997749 | 0.995962 | 0.996854 |
| | PEPPER | INDEL | 504501 | 399453 | 105048 | 54650 | 0.791778 | 0.882125 | 0.834514 |
| | | SNP | 3327495 | 3321557 | 5938 | 19046 | 0.998215 | 0.9943 | 0.996254 |
| | Clair3 | INDEL | 504501 | 401819 | 102682 | 51547 | 0.796468 | 0.88874 | 0.840078 |
| | | SNP | 3327495 | 3320762 | 6733 | 21388 | 0.997977 | 0.993603 | 0.995785 |
| 35x | DeepVariant | INDEL | 504501 | 416788 | 87713 | 55671 | 0.826139 | 0.885965 | 0.855007 |
| | | SNP | 3327495 | 3320458 | 7037 | 12739 | 0.997885 | 0.99618 | 0.997032 |
| | PEPPER | INDEL | 504501 | 407403 | 97098 | 52990 | 0.807537 | 0.88734 | 0.845559 |
| | | SNP | 3327495 | 3322620 | 4875 | 17284 | 0.998535 | 0.994826 | 0.996677 |
| | Clair3 | INDEL | 504501 | 408248 | 96253 | 47749 | 0.809211 | 0.897618 | 0.851125 |
| | | SNP | 3327495 | 3321463 | 6032 | 18754 | 0.998187 | 0.994388 | 0.996284 |

Supplementary Table 3: Oxford Nanopore Technologies variant calling performance comparison between DeepVariant, PEPPER and Clair3 at different coverages between 10x to 35x.

| Cov. | Caller | Type | Total | TP | FN | FP | Recall | Precision | F1 |
|---|---|---|---|---|---|---|---|---|---|
| 40x | DeepVariant | INDEL | 504501 | 423764 | 80737 | 52909 | 0.839967 | 0.892638 | 0.865502 |
| | | SNP | 3327495 | 3320622 | 6873 | 12133 | 0.997934 | 0.996361 | 0.997147 |
| | PEPPER | INDEL | 504501 | 414119 | 90382 | 50919 | 0.820849 | 0.892862 | 0.855342 |
| | | SNP | 3327495 | 3323047 | 4448 | 16226 | 0.998663 | 0.995142 | 0.9969 |
| | Clair3 | INDEL | 504501 | 413098 | 91403 | 44926 | 0.818825 | 0.90416 | 0.859379 |
| | | SNP | 3327495 | 3321714 | 5781 | 16813 | 0.998263 | 0.994966 | 0.996612 |
| 45x | DeepVariant | INDEL | 504501 | 429614 | 74887 | 50066 | 0.851562 | 0.899097 | 0.874684 |
| | | SNP | 3327495 | 3320801 | 6694 | 10871 | 0.997988 | 0.996738 | 0.997363 |
| | PEPPER | INDEL | 504501 | 419218 | 85283 | 49312 | 0.830956 | 0.897071 | 0.862748 |
| | | SNP | 3327495 | 3323348 | 4147 | 14599 | 0.998754 | 0.995628 | 0.997188 |
| | Clair3 | INDEL | 504501 | 416625 | 87876 | 42430 | 0.825816 | 0.90973 | 0.865744 |
| | | SNP | 3327495 | 3322293 | 5202 | 12983 | 0.998437 | 0.996109 | 0.997272 |
| 50x | DeepVariant | INDEL | 504501 | 434092 | 70409 | 47794 | 0.860438 | 0.904207 | 0.88178 |
| | | SNP | 3327495 | 3320812 | 6683 | 10474 | 0.997992 | 0.996857 | 0.997424 |
| | PEPPER | INDEL | 504501 | 423610 | 80891 | 47830 | 0.839661 | 0.90081 | 0.869161 |
| | | SNP | 3327495 | 3323553 | 3942 | 14460 | 0.998815 | 0.995669 | 0.99724 |
| | Clair3 | INDEL | 504501 | 419323 | 85178 | 40845 | 0.831164 | 0.913352 | 0.870322 |
| | | SNP | 3327495 | 3322322 | 5173 | 12292 | 0.998445 | 0.996315 | 0.997379 |
| 55x | DeepVariant | INDEL | 504501 | 438117 | 66384 | 45778 | 0.868417 | 0.908663 | 0.888084 |
| | | SNP | 3327495 | 3320863 | 6632 | 9938 | 0.998007 | 0.997017 | 0.997512 |
| | PEPPER | INDEL | 504501 | 427106 | 77395 | 46546 | 0.846591 | 0.903942 | 0.874327 |
| | | SNP | 3327495 | 3323552 | 3943 | 13473 | 0.998815 | 0.995964 | 0.997387 |
| | Clair3 | INDEL | 504501 | 421823 | 82678 | 39301 | 0.836119 | 0.91683 | 0.874616 |
| | | SNP | 3327495 | 3322252 | 5243 | 11429 | 0.998424 | 0.996573 | 0.997498 |
| 60x | DeepVariant | INDEL | 504501 | 441442 | 63059 | 43947 | 0.875007 | 0.912632 | 0.893423 |
| | | SNP | 3327495 | 3320861 | 6634 | 9573 | 0.998006 | 0.997127 | 0.997566 |
| | PEPPER | INDEL | 504501 | 430032 | 74469 | 45458 | 0.852391 | 0.906561 | 0.878642 |
| | | SNP | 3327495 | 3323667 | 3828 | 12639 | 0.99885 | 0.996213 | 0.997529 |
| | Clair3 | INDEL | 504501 | 423735 | 80766 | 37991 | 0.839909 | 0.91973 | 0.878009 |
| | | SNP | 3327495 | 3322068 | 5427 | 9920 | 0.998369 | 0.997024 | 0.997696 |
| 65x | DeepVariant | INDEL | 504501 | 444208 | 60293 | 42612 | 0.88049 | 0.915553 | 0.897679 |
| | | SNP | 3327495 | 3320812 | 6683 | 9294 | 0.997992 | 0.99721 | 0.997601 |
| | PEPPER | INDEL | 504501 | 432767 | 71734 | 43959 | 0.857812 | 0.909908 | 0.883093 |
| | | SNP | 3327495 | 3323726 | 3769 | 11211 | 0.998867 | 0.996639 | 0.997752 |
| | Clair3 | INDEL | 504501 | 424915 | 79586 | 37232 | 0.842248 | 0.921413 | 0.880054 |
| | | SNP | 3327495 | 3321510 | 5985 | 9341 | 0.998201 | 0.997197 | 0.997699 |

Supplementary Table 4: Oxford Nanopore Technologies variant calling performance comparison between DeepVariant, PEPPER and Clair3 at different coverages between 40x to 65x.

| Cov. | Platform | Type | Total | TP | FN | FP | Recall | Precision | F1 |
|---|---|---|---|---|---|---|---|---|---|
| 10x | R10.4.1 DeepVariant | INDEL | 504501 | 316320 | 188181 | 69165 | 0.626996 | 0.824024 | 0.712133 |
| | | SNP | 3327495 | 3152000 | 175495 | 56238 | 0.947259 | 0.982476 | 0.964546 |
| | R9.4.1 PEPPER | INDEL | 504501 | 209342 | 295159 | 40606 | 0.414949 | 0.839909 | 0.555472 |
| | | SNP | 3327495 | 2984659 | 342836 | 57620 | 0.896969 | 0.981062 | 0.937133 |
| 15x | R10.4.1 DeepVariant | INDEL | 504501 | 359668 | 144833 | 69022 | 0.712918 | 0.843013 | 0.772527 |
| | | SNP | 3327495 | 3288775 | 38720 | 25744 | 0.988364 | 0.992235 | 0.990296 |
| | R9.4.1 PEPPER | INDEL | 504501 | 276696 | 227805 | 50025 | 0.548455 | 0.849811 | 0.666659 |
| | | SNP | 3327495 | 3230875 | 96620 | 18904 | 0.970963 | 0.994184 | 0.982436 |
| 20x | R10.4.1 DeepVariant | INDEL | 504501 | 380627 | 123874 | 65512 | 0.754462 | 0.857262 | 0.802584 |
| | | SNP | 3327495 | 3312923 | 14572 | 18308 | 0.995621 | 0.994506 | 0.995063 |
| | R9.4.1 PEPPER | INDEL | 504501 | 311062 | 193439 | 53260 | 0.616574 | 0.856908 | 0.717141 |
| | | SNP | 3327495 | 3286111 | 41384 | 11946 | 0.987563 | 0.996379 | 0.991951 |
| 25x | R10.4.1 DeepVariant | INDEL | 504501 | 395117 | 109384 | 62184 | 0.783184 | 0.868111 | 0.823463 |
| | | SNP | 3327495 | 3318240 | 9255 | 15473 | 0.997219 | 0.99536 | 0.996289 |
| | R9.4.1 PEPPER | INDEL | 504501 | 332679 | 171822 | 52935 | 0.659422 | 0.865802 | 0.748649 |
| | | SNP | 3327495 | 3303138 | 24357 | 9843 | 0.99268 | 0.99703 | 0.99485 |
| 30x | R10.4.1 DeepVariant | INDEL | 504501 | 407454 | 97047 | 58882 | 0.807638 | 0.877687 | 0.841207 |
| | | SNP | 3327495 | 3320004 | 7491 | 13466 | 0.997749 | 0.995962 | 0.996854 |
| | R9.4.1 PEPPER | INDEL | 504501 | 348343 | 156158 | 51857 | 0.69047 | 0.873415 | 0.771242 |
| | | SNP | 3327495 | 3310086 | 17409 | 8939 | 0.994768 | 0.997307 | 0.996036 |
| 35x | R10.4.1 DeepVariant | INDEL | 504501 | 416788 | 87713 | 55671 | 0.826139 | 0.885965 | 0.855007 |
| | | SNP | 3327495 | 3320458 | 7037 | 12739 | 0.997885 | 0.99618 | 0.997032 |
| | R9.4.1 PEPPER | INDEL | 504501 | 360393 | 144108 | 50461 | 0.714355 | 0.880074 | 0.788603 |
| | | SNP | 3327495 | 3313305 | 14190 | 8230 | 0.995736 | 0.997523 | 0.996628 |

Supplementary Table 5: Oxford Nanopore Technologies variant calling performance comparison between R9.4.1 and R10.4 chemistry data at coverages between 10x and 35x.

| Coverage | Platform | Type | Total | TP | FN | FP | Recall | Precision | F1_Score |
|---|---|---|---|---|---|---|---|---|---|
| 40x | R10.4.1 DeepVariant | INDEL | 504501 | 423764 | 80737 | 52909 | 0.839967 | 0.892638 | 0.865502 |
| | | SNP | 3327495 | 3320622 | 6873 | 12133 | 0.997934 | 0.996361 | 0.997147 |
| | R9.4.1 PEPPER | INDEL | 504501 | 370622 | 133879 | 49181 | 0.734631 | 0.885651 | 0.803103 |
| | | SNP | 3327495 | 3315180 | 12315 | 7993 | 0.996299 | 0.997595 | 0.996947 |
| 45x | R10.4.1 DeepVariant | INDEL | 504501 | 429614 | 74887 | 50066 | 0.851562 | 0.899097 | 0.874684 |
| | | SNP | 3327495 | 3320801 | 6694 | 10871 | 0.997988 | 0.996738 | 0.997363 |
| | R9.4.1 PEPPER | INDEL | 504501 | 378872 | 125629 | 47879 | 0.750984 | 0.890526 | 0.814824 |
| | | SNP | 3327495 | 3316265 | 11230 | 7717 | 0.996625 | 0.997679 | 0.997152 |
| 50x | R10.4.1 DeepVariant | INDEL | 504501 | 434092 | 70409 | 47794 | 0.860438 | 0.904207 | 0.88178 |
| | | SNP | 3327495 | 3320812 | 6683 | 10474 | 0.997992 | 0.996857 | 0.997424 |
| | R9.4.1 PEPPER | INDEL | 504501 | 385925 | 118576 | 46668 | 0.764964 | 0.894782 | 0.824796 |
| | | SNP | 3327495 | 3316884 | 10611 | 7169 | 0.996811 | 0.997844 | 0.997327 |
| 55x | R10.4.1 DeepVariant | INDEL | 504501 | 438117 | 66384 | 45778 | 0.868417 | 0.908663 | 0.888084 |
| | | SNP | 3327495 | 3320863 | 6632 | 9938 | 0.998007 | 0.997017 | 0.997512 |
| | R9.4.1 PEPPER | INDEL | 504501 | 392084 | 112417 | 45404 | 0.777172 | 0.898791 | 0.833569 |
| | | SNP | 3327495 | 3317426 | 10069 | 7255 | 0.996974 | 0.997818 | 0.997396 |
| 60x | R10.4.1 DeepVariant | INDEL | 504501 | 441442 | 63059 | 43947 | 0.875007 | 0.912632 | 0.893423 |
| | | SNP | 3327495 | 3320861 | 6634 | 9573 | 0.998006 | 0.997127 | 0.997566 |
| | R9.4.1 PEPPER | INDEL | 504501 | 397171 | 107330 | 44010 | 0.787255 | 0.902738 | 0.841051 |
| | | SNP | 3327495 | 3317700 | 9795 | 7126 | 0.997056 | 0.997857 | 0.997457 |
| 65x | R10.4.1 DeepVariant | INDEL | 504501 | 444208 | 60293 | 42612 | 0.88049 | 0.915553 | 0.897679 |
| | | SNP | 3327495 | 3320812 | 6683 | 9294 | 0.997992 | 0.99721 | 0.997601 |
| | R9.4.1 PEPPER | INDEL | 504501 | 401236 | 103265 | 42583 | 0.795313 | 0.906457 | 0.847255 |
| | | SNP | 3327495 | 3317934 | 9561 | 7111 | 0.997127 | 0.997862 | 0.997494 |

Supplementary Table 6: Oxford Nanopore Technologies variant calling performance comparison between R9.4.1 and R10.4 chemistry data at coverages between 40x and 65x.

| Cov | Type | Type | Total | TP | FN | FP | Recall | Precision | F1 |
|---|---|---|---|---|---|---|---|---|---|
| 10x | R10.4.1 Duplex | INDEL | 11256 | 7610 | 3646 | 1649 | 0.676084 | 0.825207 | 0.743239 |
| | | SNP | 71333 | 67511 | 3822 | 905 | 0.94642 | 0.986779 | 0.966178 |
| | R10.4.1 Simplex | INDEL | 11256 | 6751 | 4505 | 1614 | 0.599769 | 0.810073 | 0.689236 |
| | | SNP | 71333 | 66281 | 5052 | 1336 | 0.929177 | 0.980252 | 0.954031 |
| 15x | R10.4.1 Duplex | INDEL | 11256 | 8624 | 2632 | 1529 | 0.766169 | 0.853051 | 0.807279 |
| | | SNP | 71333 | 70666 | 667 | 200 | 0.990649 | 0.997179 | 0.993904 |
| | R10.4.1 Simplex | INDEL | 11256 | 7842 | 3414 | 1572 | 0.696695 | 0.836454 | 0.760205 |
| | | SNP | 71333 | 70054 | 1279 | 533 | 0.98207 | 0.992453 | 0.987234 |
| 20x | R10.4.1 Duplex | INDEL | 11256 | 9092 | 2164 | 1383 | 0.807747 | 0.871397 | 0.838365 |
| | | SNP | 71333 | 71184 | 149 | 113 | 0.997911 | 0.998416 | 0.998164 |
| | R10.4.1 Simplex | INDEL | 11256 | 8353 | 2903 | 1498 | 0.742093 | 0.851566 | 0.793069 |
| | | SNP | 71333 | 70967 | 366 | 223 | 0.994869 | 0.996869 | 0.995868 |
| 25x | R10.4.1 Duplex | INDEL | 11256 | 9359 | 1897 | 1312 | 0.831468 | 0.880488 | 0.855276 |
| | | SNP | 71333 | 71250 | 83 | 93 | 0.998836 | 0.998697 | 0.998767 |
| | R10.4.1 Simplex | INDEL | 11256 | 8690 | 2566 | 1425 | 0.772033 | 0.862849 | 0.814918 |
| | | SNP | 71333 | 71169 | 164 | 162 | 0.997701 | 0.99773 | 0.997716 |
| 30x | R10.4.1 Duplex | INDEL | 11256 | 9528 | 1728 | 1282 | 0.846482 | 0.884868 | 0.865249 |
| | | SNP | 71333 | 71269 | 64 | 79 | 0.999103 | 0.998893 | 0.998998 |
| | R10.4.1 Simplex | INDEL | 11256 | 8974 | 2282 | 1369 | 0.797264 | 0.871335 | 0.832655 |
| | | SNP | 71333 | 71211 | 122 | 141 | 0.99829 | 0.998025 | 0.998157 |
| 35x | R10.4.1 Duplex | INDEL | 11256 | 9700 | 1556 | 1162 | 0.861763 | 0.896453 | 0.878766 |
| | | SNP | 71333 | 71276 | 57 | 78 | 0.999201 | 0.998907 | 0.999054 |
| | R10.4.1 Simplex | INDEL | 11256 | 9145 | 2111 | 1338 | 0.812456 | 0.876088 | 0.843073 |
| | | SNP | 71333 | 71236 | 97 | 123 | 0.99864 | 0.998277 | 0.998459 |
| 40x | R10.4.1 Duplex | INDEL | 11256 | 9830 | 1426 | 1068 | 0.873312 | 0.905109 | 0.888926 |
| | | SNP | 71333 | 71279 | 54 | 73 | 0.999243 | 0.998978 | 0.99911 |
| | R10.4.1 Simplex | INDEL | 11256 | 9318 | 1938 | 1204 | 0.827825 | 0.888961 | 0.857304 |
| | | SNP | 71333 | 71233 | 100 | 123 | 0.998598 | 0.998277 | 0.998438 |
| 45x | R10.4.1 Duplex | INDEL | 11256 | 9920 | 1336 | 1011 | 0.881308 | 0.910523 | 0.895677 |
| | | SNP | 71333 | 71283 | 50 | 69 | 0.999299 | 0.999033 | 0.999166 |
| | R10.4.1 Simplex | INDEL | 11256 | 9443 | 1813 | 1171 | 0.83893 | 0.892834 | 0.865043 |
| | | SNP | 71333 | 71239 | 94 | 127 | 0.998682 | 0.998222 | 0.998452 |
| 50x | R10.4.1 Duplex | INDEL | 11256 | 10024 | 1232 | 951 | 0.890547 | 0.916248 | 0.903215 |
| | | SNP | 71333 | 71286 | 47 | 69 | 0.999341 | 0.999034 | 0.999187 |
| | R10.4.1 Simplex | INDEL | 11256 | 9530 | 1726 | 1108 | 0.84666 | 0.898914 | 0.872005 |
| | | SNP | 71333 | 71249 | 84 | 120 | 0.998822 | 0.99832 | 0.998571 |

Supplementary Table 7: Oxford Nanopore Technologies variant calling performance comparison between Simplex and Duplex data types.
